# Multiplexed changes in synaptic transmission underlie stress-induced reduction of persistent firing in the parietal cortex

**DOI:** 10.1101/2025.08.27.672720

**Authors:** Archana Proddutur, Daniel Jun Rindner, Ghalia Azouz, Kevin Beier, Gyorgy Lur

## Abstract

Repeated exposure to stress disrupts cognitive processes, including attention and working memory. A key mechanism supporting these functions is the ability of neurons to sustain action potential firing, even after a stimulus is no longer present. How stress impacts this persistent neuronal activity is currently unknown. We found that repeated exposure to multiple concurrent stressors during adolescence (aRMS) impedes the ability of layer 5 pyramidal neurons (L5 PNs) in the posterior parietal cortex (PPC) to produce persistent firing. To determine the mechanisms underlying this effect, we complemented computational modelling with whole-cell patch clamp electrophysiology in acute brain slices from male mice. Our model predicted that altered intrinsic excitability, reduced local connectivity, diminished glutamatergic transmission, or enhanced inhibition could explain diminished persistent activity. In *ex vivo* experiments, we found minimal effect of aRMS on excitability and recurrent connectivity. However, stress exposure altered the properties of excitatory connections between L5 PNs, specifically affecting decay kinetics and short-term synaptic dynamics. In addition, aRMS increased inhibitory tone in the PPC, altering both GABAa and GABAb receptor-mediated responses. Incorporating the observed physiological changes into our network model, we found that no single parameter was sufficient alone to reproduce the stress-induced reduction in persistent firing. Rather, a combination of altered excitatory and inhibitory synaptic transmission was necessary to impact sustained activity. These data suggest that a multitude of converging changes in neural and circuit function underpin the effects of stress on cognitive processes.

**Highlights:**

- Stress exposure impairs persistent firing in the posterior parietal cortex.
- GABAergic inhibition is enhanced following stress.
- Stress alters excitatory transmission between layer 5 pyramidal cells.
- Combined changes in inhibition and excitation impact persistent firing.

## Introduction

Chronic stress is a well-established risk factor for cognitive dysfunction, frequently impacting attention, decision-making, as well as short-, and long-term memory (McEwen 2017, Arnsten 2009, McEwen, Nasca, and Gray 2016, Sousa and Almeida 2012, Park and Lur 2024, Libovner et al. 2020). These higher-order processes rely on working memory, the capacity of neural circuits to maintain information for short periods of time, even in the absence of continuous sensory input (Goldman-Rakic 1995, Gold and Shadlen 2007). The mechanism underlying these processes is thought to be persistent or sustained action potential firing in response to transient stimuli, that enables the temporary retention of cognitive representations (Fuster and Alexander 1971, Larimer and Strowbridge 2010, Wang et al. 2013, Zylberberg and Strowbridge 2017). This form of sustained activity has been reported in the hippocampus, parietal regions, and the prefrontal cortex (Andersen and Cui 2009, Fuster 2021, Shadlen and Newsome 2001).

Substantial evidence suggests that stress exposure can significantly disrupt the function of neuronal circuits implicated in persistent activity. For example, in the prefrontal cortex and hippocampus, chronic restraint stress has been found to modify synaptic connectivity, decrease plasticity, and impair cognitive functions (McEwen, Nasca, and Gray 2016, Shonkoff et al. 2012, Sousa and Almeida 2012, Sandi and Haller 2015). Repeated stress exposure during early- to mid-adolescence altered the connectivity of posterior parietal cortex (PPC) and degraded visuospatial working memory in mice (Libovner et al. 2020). However, the effects of stress on persistent action potential firing remain unexplored.

Theoretical work indicates that cortical networks with recurrent excitatory connectivity can amplify input-specific signals, supporting sustained action potential firing (Compte et al. 2000, Wang 2008, Peron et al. 2020). In such networks, N-methyl-D-aspartate-type glutamate receptors (NMDARs) enable network-level persistence due to their long decay kinetics (Wang 1999). This high-gain excitatory framework necessitates precise inhibitory control to prevent runaway excitation, typically mediated by gamma-aminobutyric acid (GABA)ergic interneurons (Najafi et al. 2020, Papoutsi, Sidiropoulou, and Poirazi 2014, Sadeh and Clopath 2021, Destexhe, Contreras, and Steriade 1998, Garcia Del Molino et al. 2017). Indeed, disruption of the excitatory-inhibitory balance has been linked to cognitive impairments in animal models (Page and Coutellier 2019, Rodrigues et al. 2024, Wang et al. 2019). While there is evidence that stress can alter both intrinsic excitability and synaptic connectivity of cortical neurons (Nawreen, Baccei, and Herman 2021, Urban and Valentino 2017), how these inherently intertwined changes might impact the mechanisms underpinning persistent activity is unknown.

The current study is based on an observation that repeated exposure to multiple concurrent stressors in adolescent mice (aRMS for short) diminished the ability of layer 5 pyramidal neurons (L5 PNs) in the PPC to produce persistent firing. To identify the mechanisms underlying this effect, we utilized a computational model of a persistent firing network (Papoutsi, Sidiropoulou, and Poirazi 2014). The model predicted that changes in the number or strength of excitatory and inhibitory synaptic connections, as well as altered intrinsic properties, could explain the observed reduction in persistent firing. We evaluated these model predictions using whole-cell patch clamp electrophysiology in acute brain slices and found that aRMS affected both excitatory and inhibitory synaptic transmission. To assess how each change could impact persistent firing, we systematically explored their separate and collective consequences in our computational model. These tests indicated that while individually unremarkable, the combination of stress effects on cellular and synaptic function can substantially impact persistent firing.

## Results

### Repeated exposure to multiple concurrent stresses during adolescence reduced persistent firing in L5 PNs of the PPC

Previous data indicate that repeated exposure to multiple concurrent stresses in early- to mid-adolescence (aRMS) alters the synaptic connectivity of the posterior parietal cortex (PPC) and diminishes visuo-spatial working memory in mice (Libovner et al. 2020). Based on these prior results, we hypothesized that aRMS impedes the ability of the PPC circuit to produce sustained neuronal activity. To test this hypothesis, we made whole cell patch clamp recordings from layer 5 pyramidal cells (L5 PNs) in acute brain slices containing the PPC. To promote persistent activity, we used aCSF containing 1.2 mM Ca^2+^ and 10 µM carbachol (see Methods). Somatic current injections (ranging from 0.25 – 4 seconds) evoked action potential firing from the recorded neurons, which was followed by a period of sustained firing, observable in both control (Figure 1A) and aRMS (Figure 1B) mice. However, the duration of persistent activity in response to the same set of stimuli was markedly shorter in aRMS animals (fixed effect of stress: F_1, 152_ = 13.42, p = 0.0003, n_ctr_ = 20, n_aRMS_ = 22 cells, repeated measures mixed effects model, Figure 1C). To estimate the length of the persistent activity produced with different stimulus lengths, we fitted a one-phase association curve to the data. This showed that persistent firing duration plateaued around 33.9 s in controls, compared to 11.6 s in aRMS (F_2, 158_ = 8.37, p = 0.0003, sum-of-squares F test). Similarly, the frequency of action potentials, calculated over the first 10 seconds of persistent firing, was lower in aRMS mice than in controls (fixed effect of stress: F_1, 40_ = 5.97, p = 0.019, repeated measures mixed effects model, Figure 1D). Fitting the firing frequency data with a one-phase association curve indicated that on average, persistent firing frequency plateaued at 9.4 Hz in controls and 3.7 Hz in aRMS (F_2, 91_ = 9.04, p = 0.0003, sum-of-squares F test). We found no difference in the proportion of control and aRMS neurons that were able to produce sustained activity (p = 0.7585, Fisher’s exact test, Figure 1E), or in the action potential firing rates during the initial current injection step (fixed effect of stress: F_1, 53_ = 1.787, p = 0.187, repeated measures mixed effects model, SFigure 1A). In addition, L5 PNs from control and aRMS mice produced indistinguishable afterdepolarization following a 2 s current injection when measured at the membrane potential hyperpolarized to −75mV (control: 1.99 ± 0.85, n = 15, aRMS: 1.37 ± 1.58, n = 10, p = 0.45, Welch’s test, SFigure 1B).

**Figure 1:**
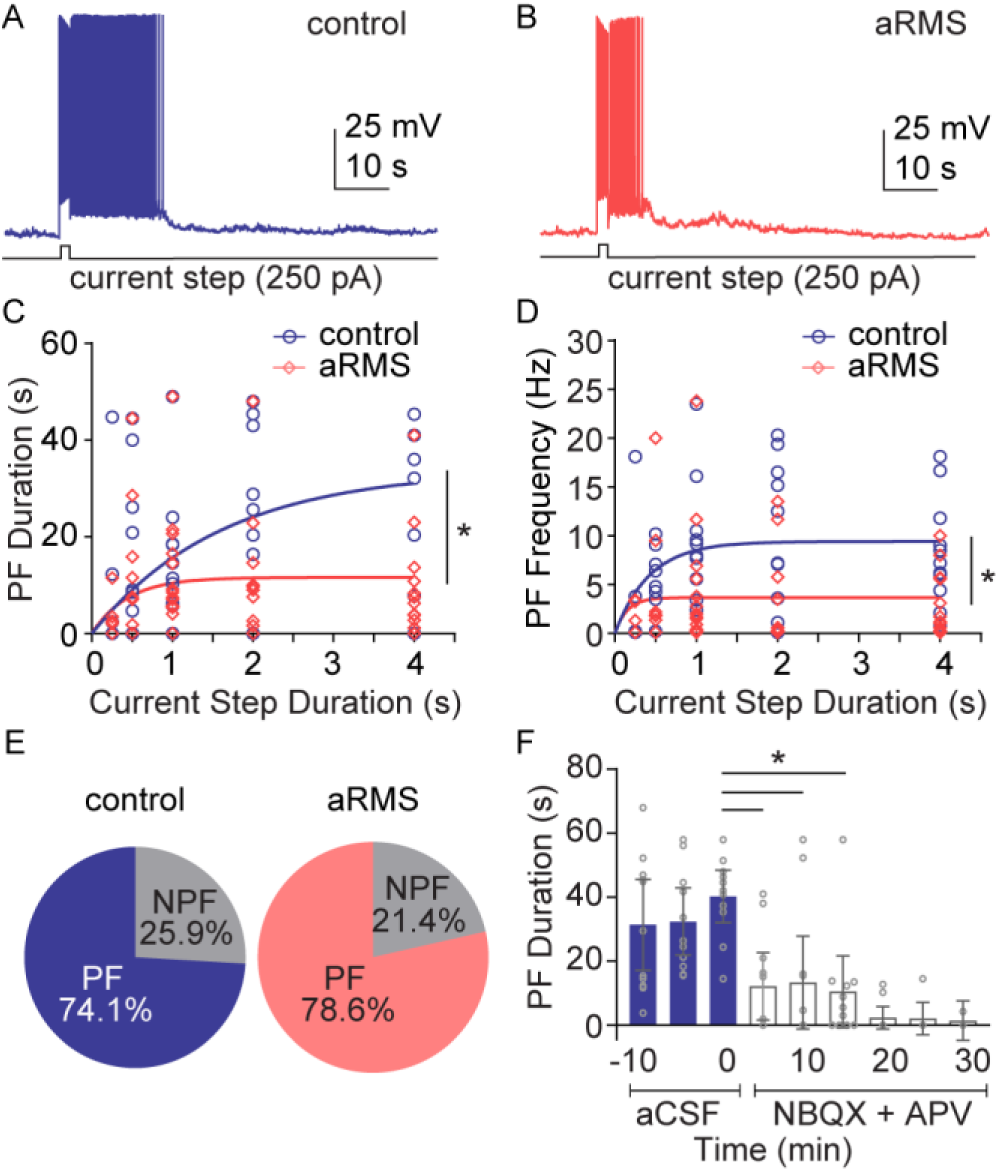
aRMS reduces persistent activity in L5 PNs of the PPC. (**A**) Representative voltage trace showing persistent activity in control male mice (blue). (**B**) Representative trace of persistent activity in aRMS exposed animals. (**C**) Summary plot showing the duration of persistent activity in response to increasing current step length in control (blue) and aRMS (salmon) mice. (**D**) Frequency of persistent activity during the first 10 seconds after induction in control (blue)and aRMS (salmon) mice. Overlaid curves in C and D represent exponential one-phase association nonlinear fits. (**E**) Pie charts show the proportion of cells that produced persistent (PF) compared to those that did not (NPF) in control and aRMS groups. (**F**) Persistent firing duration in controls, recorded every 5 minutes before and after glutamate blockers (NBQX: 10 µM and AP5: 100 µM). Bars indicate mean ± 95 CI; *: p < 0.05.

Most theoretical frameworks argue that persistent firing relies on a recurrent excitatory network (Compte et al. 2000, Wang 1999, Amit and Brunel 1997, Tegner, Compte, and Wang 2002). This suggests that blocking glutamate receptors would eliminate persistent firing in our slice preparation. To test this, we recorded persistent activity in L5 PNs in the PPC for three consecutive sweeps with 5-minute intervals to establish a baseline of sustained firing duration, then bath applied a cocktail of antagonists for AMPA and NMDA type glutamate receptors (10 μM NBQX and 50 μM AP5). We measured persistent firing duration every 5 minutes after drug flow-in and observed rapid reduction in sustained activity following glutamate receptor block (persistent firing duration in aCSF: 31.37 ± 14.2 s; in the first 15 minutes in glutamate blockers: 10.46± 11.2 s; n = 11; fixed effect of blockers: F_3, 21.6_ = 7.4, p = 0.0012, mixed-effects analysis, Figure 1F). Post-hoc tests corrected for multiple comparisons using Sidak’s method indicated a significant reduction in persistent firing duration in the first 5 minutes after glutamate receptor block (p_5min_ = 0.0035) that was maintained during later time points (p_10min_ = 0.017, p_15min_ = 0.004). Persistent firing duration practically went to zero after 20 minutes in glutamate block. Without the presence of glutamate receptor antagonists, persistent activity remained stable (persistent firing duration at the start: 42.7 ± 16.02 s and after 30 minutes: 45.3 ± 29.8 s, fixed effect of time: F_4, 16_ = 0.497, p = 0.74, n = 5, one-way repeated measures ANOVA, SFigure 1C).

These findings indicate that aRMS exposure diminishes the ability of L5 PNs to produce persistent firing. In control animals, sustained activity appears to rely on excitatory neurotransmission.

### Recurrent network model predicts synaptic and circuit mechanisms underlying stress-induced reduction in persistent activity

The ability of a cortical circuit to produce persistent firing could be impacted by a plethora of variables. To determine what cellular and circuit mechanisms could contribute to the aRMS effect on persistent firing, we employed a computational approach. We adapted previously published (see methods) small model circuit consisting of seven, recurrently connected pyramidal cells and two interneurons. The network was stimulated by excitatory input of varying strength and duration, and we recorded the spiking output of each cell (Figure 2A). In this model, we can independently vary parameters and generate predictions of what stress-induced changes would result in reduced persistent firing. As expected, increasing the stimulus duration or the weight of the NMDA component in the synaptic input driving the network increased the probability of evoking persistent firing (Figure 2B). For a systematic exploration of other variables, we fixed these two parameters at values that produced approximately 60% chance of generating sustained activity (2 s duration and 0.35 NMDA weight, see Methods). In terms of external factors, we found that increasing the decay time constant of NMDA-receptors in the synaptic drive of the network will increase persistent firing probability. With regards to variables within the recurrent network, decreasing the number of synapses between pyramidal neurons or decreasing the strength of these recurrent connections is predicted to reduce the probability of persistent firing. In addition, diminished persistent firing may be caused by increasing inhibitory synaptic conductance through GABAa-, or GABAb-receptors. As for intrinsic properties, the model predicted that changing the input resistance of participating pyramidal neurons will have a marked effect on the network’s ability to produce persistent firing. Conversely, altering the h-current appeared to have a lesser effect. In the following experiments, we use *ex vivo* physiology and histology methods to test the above model predictions, comparing cellular and circuit properties between control and aRMS mice.

**Figure 2:**
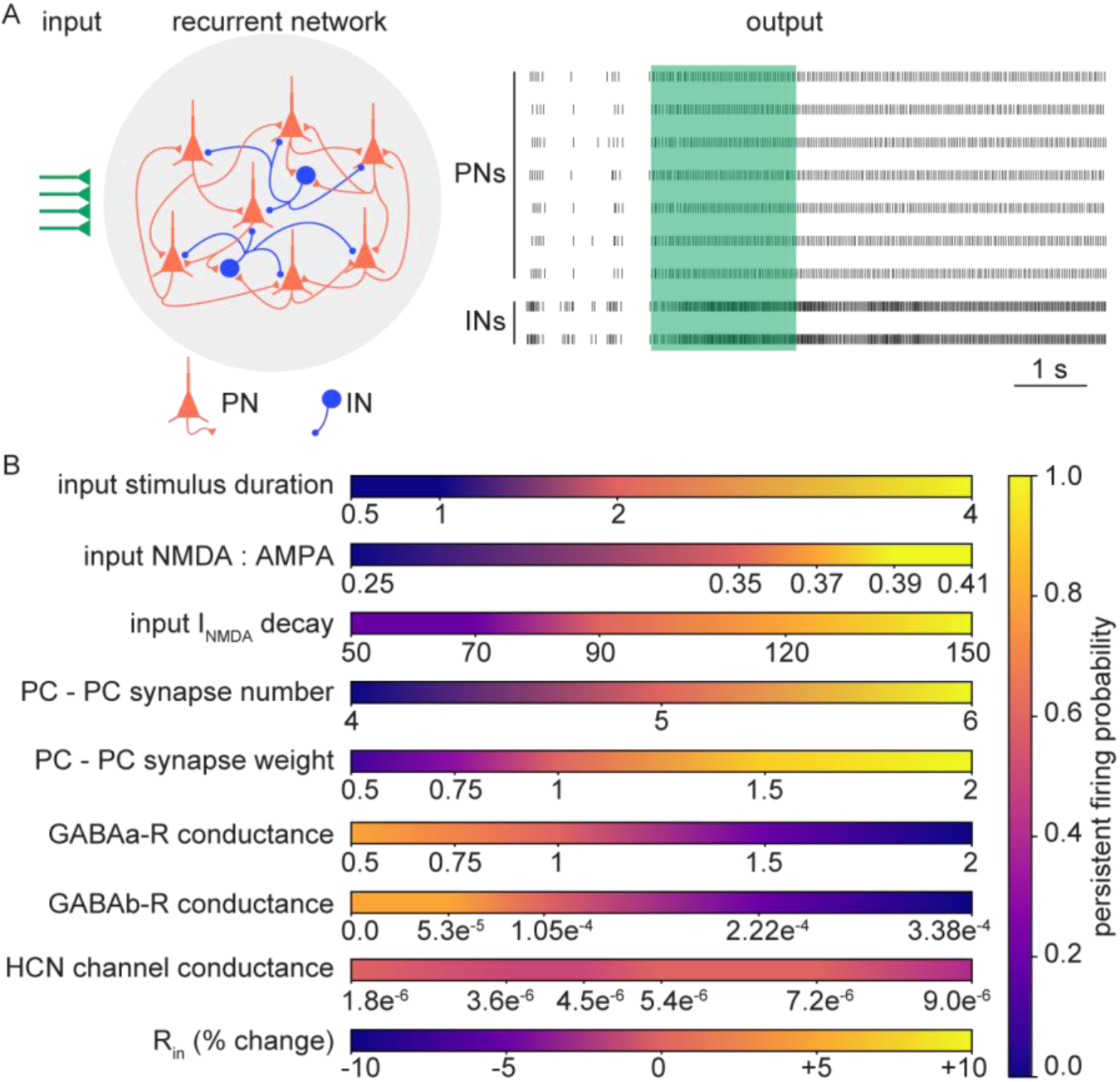
Computational model predicts potential synaptic and circuit mechanisms underlying stress effect on persistent firing. (**A**) Left: Schematic of *in silico* neuronal network containing 7 pyramidal neurons (PNs, orange) and 2 interneurons (INs, blue). Green arrows represent external synaptic drive. Right: Representative spike raster plot showing action potentials in PNs and INs. The green shaded area indicates the time when external synaptic drive was applied. (**B**) Heatmaps show persistent firing probabilities in response to varied synaptic or ionic channel parameters while all other variables were kept constant. Warmer colors indicate higher persistent firing probability.

### aRMS reduces input resistance of L5 PNs in the PPC

To assess aRMS effects on the intrinsic properties of L5 PNs of the PPC, we used whole-cell patch clamp recordings to measure responses to a series of hyperpolarizing and depolarizing current pulses in regular aCSF extracellular solution (Figure 3A, B). Both groups showed increasing action potential firing in response to depolarizing pulses, and we found the input-output relationship unaffected by aRMS (main effect of stress: F_1,56_ = 2.23, p = 0.14, main effect of current step: F_2.5, 139.6_ = 264.8, p < 0.0001, interaction: F_8, 448_ = 0.8, p = 0.6, n_ctr_ = 36, n_aRMS_ = 22, two-way repeated measures ANOVA, Figure 3C). Action potential threshold (control: −41.59 ± 1.94, n = 36, aRMS: −40.52 ± 2.47, n = 22, p = 0.48, Welch’s t-test, Figure 3D, E) and sag amplitude (control: 2.1 ± 0.35 mV, n = 36, aRMS: 1.85 ± 0.43 mV, n = 22, p = 0.43, Welch’s t-test, Figure 3F) also appeared unaltered by aRMS. In contrast, we found that input resistance of L5 PNs from aRMS animals was lower than from controls (tested at positive step current (50pA) with no overriding action potentials: control: 195.7 ± 18.7 MΩ, n = 36; aRMS: 160 ± 21.1 MΩ, n = 22 p = 0.012, Welch’s t-test, Figure 3G).

**Figure 3:**
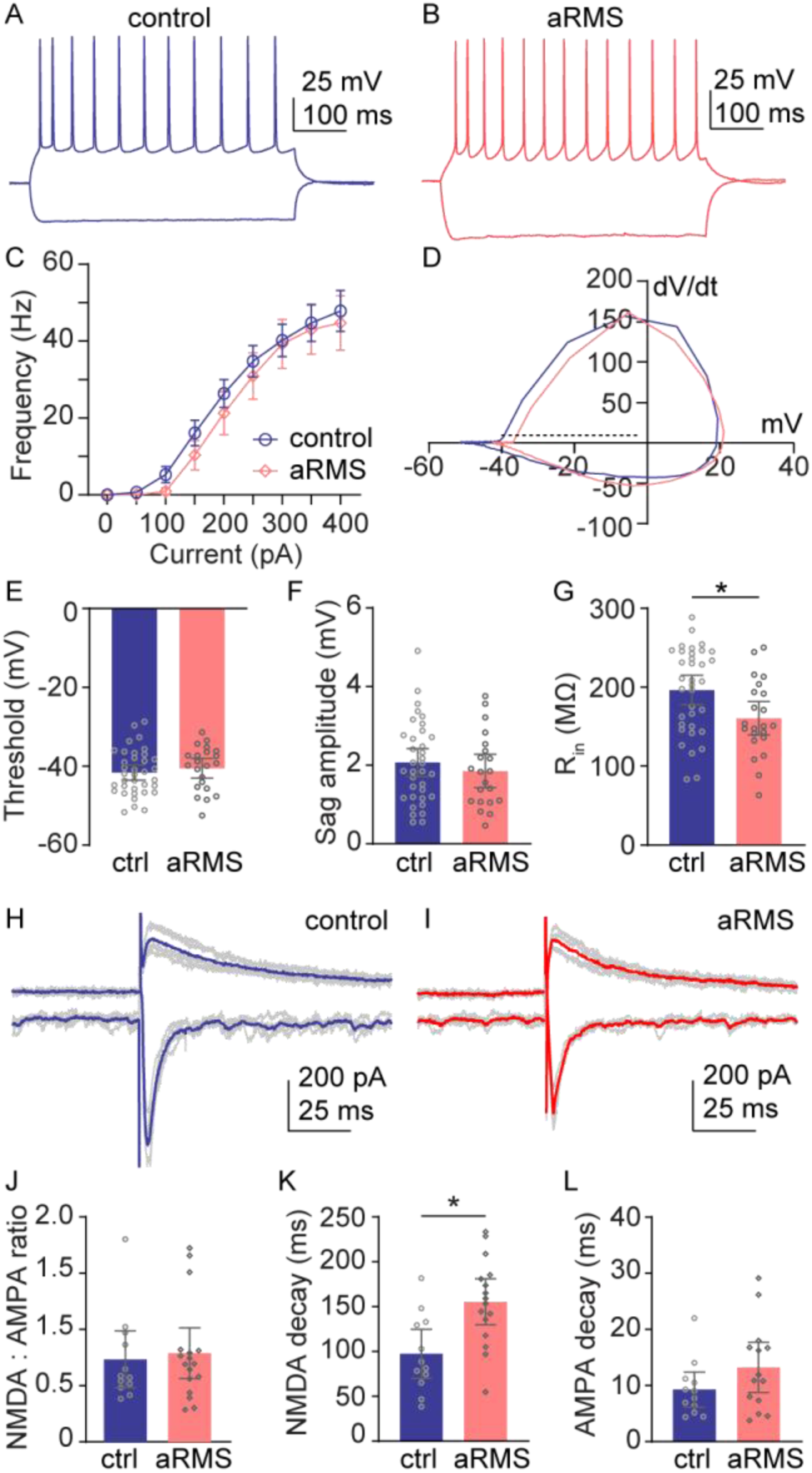
Effect of aRMS on intrinsic properties and overall synaptic drive of L5 pyramidal cells in PPC. (**A**) Example membrane voltage responses to hyperpolarizing and depolarizing current injections in control (blue) L5 PNs. (**B**) Example membrane potential traces from aRMS (salmon) mice. (**C**) Summary graph of the firing frequency - current input relation in control (blue) and aRMS (salmon) mice. (**D**) An example phase plot showing the first time derivative (dV/dt) of the voltage trace a control (blue) and an aRMS (salmon) cell. The dashed line represents the 10 mV/ms threshold. (**E**) Population data showing action potential threshold. (**F**) Population data of SAG amplitude. (**G**) Population data of input resistance measured at 50 pA depolarizing current step. (**H**) Representative AMPA and NMDA receptor currents evoked by electrical stimulation in control (blue) mice. (**I**) Representative AMPA and NMDA receptor currents in recorded in aRMS exposed animals (salmon). (**J**) Population data of NMDA : AMPA ratio in control (blue) and aRMS (salmon). (**K**) Population data of NMDA-receptor current decay kinetics of NMDA currents in control (blue) and aRMS (salmon) mice. (**L**) Population data of AMPA-receptor currents in control (blue) and aRMS (salmon) exposed animals. Bars indicate mean ± 95 CI; *: p < 0.05.

Overall, in these experiments, aRMS did not alter the input/output curve of L5 PNs, despite a detectable reduction in input resistance in stressed animals.

### aRMS only affected evoked currents through NMDA-, not AMPA-receptors in L5 PNs

To test the effect of aRMS on glutamatergic drive to the PPC circuit, we used a bipolar stimulating electrode to evoke excitatory currents, recorded in voltage clamp. We observed robust AMPA-, and NMDA-receptor driven responses in L5 PNs from both control and aRMS animals (Figure 3H, I). However, we did not detect differences in the NMDA : AMPA ratio (control: 0.73 ± 0.25, n = 12, aRMS: 0.79 ± 0.22, n = 17, p = 0.73, unpaired t-test, Figure 3J). Next, we measured the rise and decay kinetics for each glutamate receptor subtype. In aRMS mice, we measured markedly slower NMDA-receptor decay than in controls (control: 97.46 ± 27.26 ms, n = 12, aRMS:155.4 ± 25.6 ms, n=17, p = 0.0028, unpaired t-test, Figure 3K) with no change in rise times (SFigure 2A). In contrast, AMPA-receptor decay (control: 9.27 ± 3.14 ms, n = 12, aRMS: 13.24 ± 4.48 ms, n = 17, p = 0.14, unpaired t-test, Figure 3L) and rise (control: 1.65 ± 0.82 ms, n = 12, aRMS: 1.51 ± 0.52 ms, n = 17, p = 0.74, unpaired t-test, SFigure 2B), showed no differences between groups. Finally, short-term dynamics, measured by the paired pulse ratio, remained unaffected by aRMS for both AMPA-(control: 1.09 ± 0.3, n =12, aRMS: 0.98 ± 0.11, n= 17, p = 0.396, unpaired t-test) and NMDA-receptor mediated responses (control: 0.88 ± 0.28, n =12, aRMS: 0.88 ±0.1, n = 17, p = 0.97, unpaired t-test, SFigure 2C).

According to our computational model, the observed increase in the decay constant of NMDA-receptor mediated currents is expected to lengthen persistent firing duration. Since this contrasts with what we observed in aRMS mice (see Figure 1), we expect that this change is compensated elsewhere.

### aRMS did not impact the number of local recurrent connections in PPC

Previous studies suggest that local recurrent connections, particularly those between pyramidal neurons, contribute to the emergence of persistent activity in the CA3 region of the hippocampus and in the entorhinal cortex (Cui et al. 2022, Li et al. 2024, Wang 2001). Based on this, we hypothesized that aRMS could reduce persistent firing by eliminating recurrent synapses. To test this hypothesis, we performed rabies virus (RABV) based input mapping (Beier 2022). We employed an intersectional viral strategy to specifically target L5 PNs in the PPC immediately after aRMS exposure (Figure 4A, B). To estimate the number of recurrent connections in the PPC, we calculated the average number of inputs to a starter cell. We found that the number of presynaptic neurons per starter cell was indistinguishable between control and aRMS mice (control: 3.9 ± 1.4, aRMS: 4.0 ± 1.2, p = 0.89, n = 15 slices from 5 mice each, unpaired t-test, Figure 4C). The numbers of helper construct expressing (mCherry positive: control: 204.6 ± 47.4, aRMS: 172.1 ± 34.6, n = 15 slices from 5 mice each, p = 0.24, unpaired t-test, SFigure 3A, C) neurons, RABV labeled presynaptic cells (GFP positive: control: 74.6±11.8, aRMS: 95.07±25.07, n = 15 slices from 5 mice each, p = 0.12, unpaired t-test, SFigure 3B, C), or postsynaptic starter cells (neurons co-expressing mCherry and GFP: control: 31.07 ± 9.7, aRMS: 29.73 ± 10.2, n = 15 slices from 5 mice each, p = 0.84, unpaired t-test, SFigure 3C) were also the same across conditions. Finally, we found no stress effect on the proportion of helper cells that ended up as starter neurons (control: 15.3 ± 2.5 cells, aRMS: 19.7 ± 7.9 cells, p = 0.27, Welch’s t-test, Figure 4D). These controls indicate that there were no differences in labeling efficacy between the groups.

**Figure 4:**
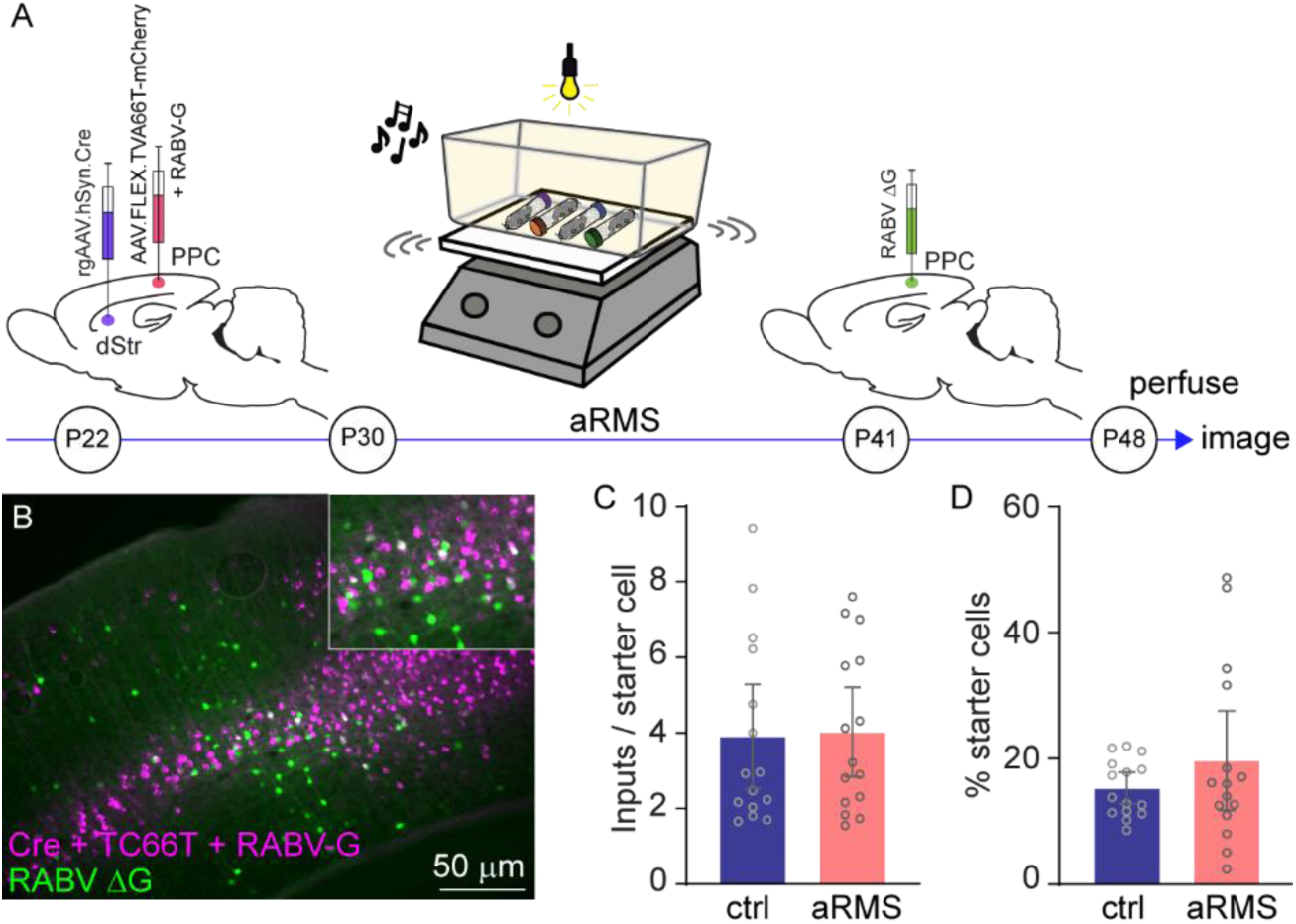
The number of local monosynaptic inputs to L5 PNs in the PPC is unaffected by aRMS. (**A**) Schematic timeline of transsynaptic tracing experiments. (**B**) example micrograph showing helper construct expressing cells in magenta and RABV expressing cells in green in the PPC. Inset: zoomed in image highlighting some co-labeled postsynaptic starter cells (white). (**C**) Population data showing the average number of inputs received by a starter cell in control (blue) and aRMS exposed (salmon) mice. (**D**) Starter neurons expressed as a proportion of all helper construct expressing cells. Bars represent population mean ± 95 CI.

The above data suggest that aRMS did not alter the number of local connections between PPC pyramidal neurons. However, this RABV-based approach only quantifies the connectivity in a unidirectional manner and does not measure synaptic strength.

### aRMS altered pre- and postsynaptic aspects of excitatory transmission in the PPC recurrent network

To expand on our histology-based connectivity measures, we used paired patch clamp recordings (Figure 5A). Dual whole cell recordings targeting two nearby pyramidal cells yielded synaptically connected pairs where action potentials evoked in one neuron resulted in postsynaptic currents in the other (Figure 5B, C). We detected no difference in the probability of PN-to-PN connectivity between control and aRMS mice (one-way connection probability in control: 19. 6%, aRMS 19. 35%; reciprocal connections in controls: 3.3%, in aRMS: 12.9%, n_ctr_ = 31, n_aRMS_ = 18, p = 0.251, Fisher’s exact test, Figure 5D). In synaptically coupled pyramidal cell pairs, we quantified response amplitude and kinetics. We did not find a statistically significant difference in the amplitude of the first response (control: 15.06 ± 2.9.3 mV, n = 15 cells, aRMS: 19.3 ± 3.4 mV, n = 12 cells, p = 0.054, Welch’s t-test, Figure 5E), or rise time (control: 2.2 ± 0.43 ms, aRMS:1.96 ± 0.8 ms, p = 0.59, Welch’s t-test, Figure 5F) between the groups. However, the decay of the synaptic response was markedly shorter in pairs from aRMS mice (control: 7.175 ± 1.5 ms, aRMS: 5.24 ± 0.8 ms, p = 0.023, Welch’s t-test, Figure 5G).

**Figure 5:**
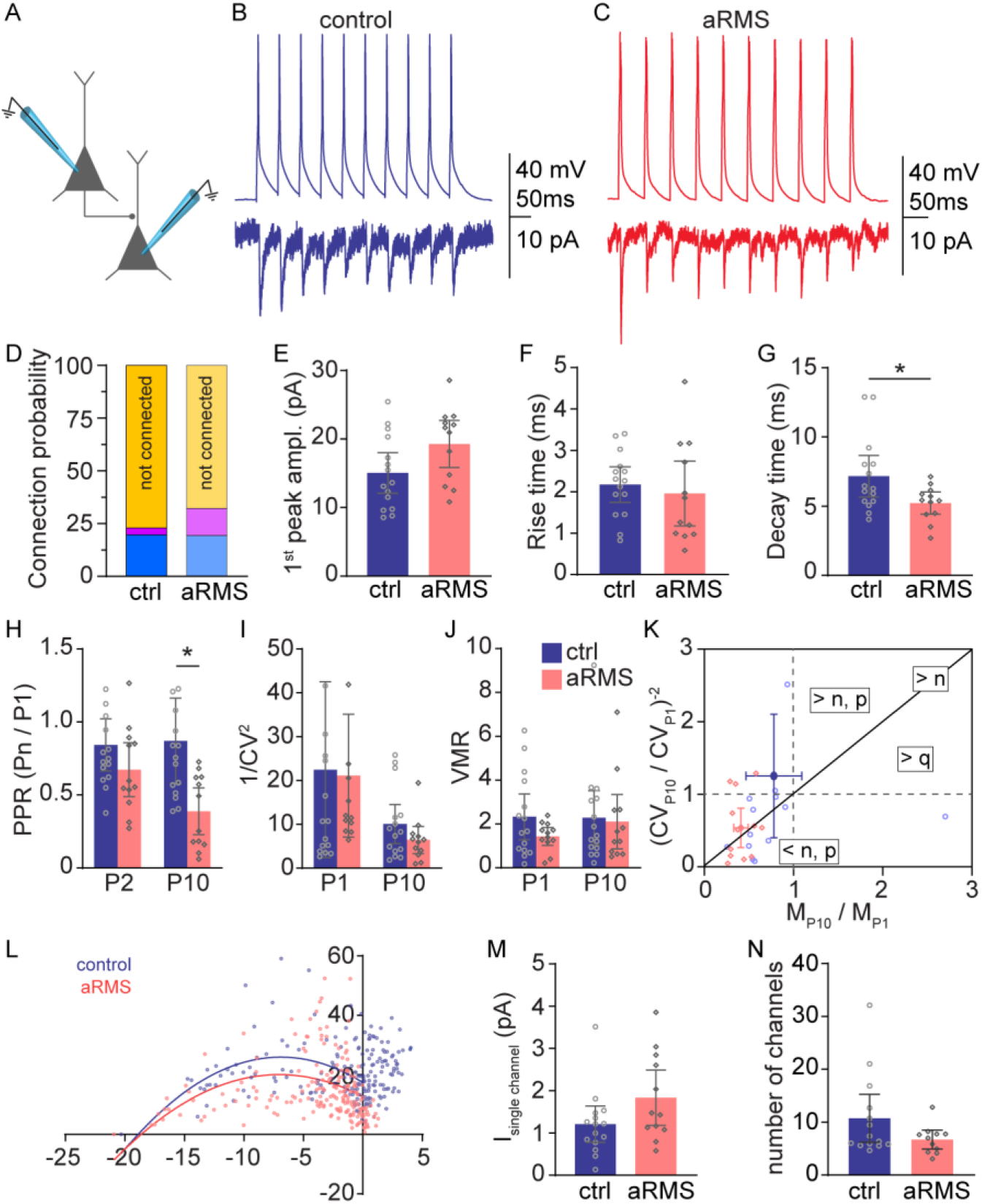
aRMS alters pre-, and postsynaptic aspects of excitatory transmission between L5 PNs in the PPC. (**A**) Schematic of paired records. (**B**) Representative traces of presynaptic action potentials (top) and postsynaptic EPSCs (bottom) recorded in synaptically coupled L5 PNs in control (blue) mice. (**C**) Representative traces from synaptically coupled L5 PNs in aRMS exposed (salmon) mice. (**D**) Connection probability in control (bold colors) and aRMS (muted colors) mice; blue hues show the proportion of unidirectional connections, magentas show bidirectional coupling, yellows indicate the proportion of cells not connected. (**E**) Population data showing the amplitude of the first response peak in coupled L5 PNs in control (blue) and in aRMS (salmon) mice. (**F**) Population data showing the rise time of EPSCs in coupled cells. (**G**) Population data showing the synaptic decay time from paired cells from control (blue) and aRMS exposed (salmon) mice. (**H**) Paired pulse ratio measured between the first and the second (P2) or tenth (P10) EPSC peak in control (blue) and aRMS (salmon) mice. (**I**) Population data of 1/CV^2^ calculated on the first (P1) and last (P10) EPSCs evoked in L5 PN pairs from control (blue) and aRMS (salmon) mice. (**J**) Population data of VMR calculated on the first (P1) and last (P10) EPSCs evoked in L5 PN pairs from control (blue) and aRMS (salmon) mice. (**K**) Topographic representation of quantal analysis calculated on the first (P1) and last (P10) EPSCs from control (blue) and aRMS (salmon) mice. Overlaid crosshairs represent the mean ± 95 CI of the population data. (**L**) Mean - variance current plot of the control (blue) and aRMS (salmon) EPSCs evoked in paired L5 PNs. Solid lines indicate fitted values (see Methods). (**M**) Estimated single channel current in control (bleu) and aRMS (salmon). (**N**) Estimated number of channels open at peak EPSC between connected pairs in control (bleu) and aRMS (salmon) mice. Bars represent mean ± 95 CI; *: p < 0.05.

To determine whether aRMS altered the properties of synaptic connections between L5 PNs, we measured the paired pulse ratio (PPR) at different time points across a 10-action potential presynaptic train. A two-way mixed effects model detected a significant effect of stress (F_1, 25_ = 9.286, p = 0.0054, n_ctr_ = 15, n_aRMS_ = 12, Figure 5H) when comparing PPR values between early and late activations. Post-hoc tests adjusted for multiple comparisons revealed no difference in the PPR between early events (p = 0.24) but showed a significant effect on the PPR between the first and last activation (p = 0.0017, Figure 5H). To determine the mechanisms underlying this difference, we applied quantal analysis to the first and last EPSCs in the response train. We calculated 1/CV^2^, a measure dependent on the number of active release sites (*n*) and release probability (*p*), but not quantal content (*q*) (Malinow and Tsien 1990, van Huijstee and Kessels 2020). Two-way mixed effects analysis showed a fixed effect for EPSC time in the train (F_1, 25_ = 5.129, p = 0.032) but not stress (F_1, 25_ = 0.1591, p = 0.69, n_ctr_ = 15, n_aRMS_ = 12, Figure 5I). Next, we assessed VMR which is dependent on *p* and *q*, but not *n* (Lupica, Proctor, and Dunwiddie 1992, van Huijstee and Kessels 2020), and found no effect of EPSC time in the train (F_1, 25_ = 0.3985, p = 0.53) or aRMS (F_1, 25_ = 1.124, p = 0.3, two-way mixed effects model, Figure 5J). Together, these two metrics suggest that the change in PPR shown in Figure 5H is likely presynaptic in origin (Kloc and Maffei 2014, van Huijstee and Kessels 2020, Wang, Fontanini, and Maffei 2012). Topographic representation (Bekkers and Stevens 1990, Malinow and Tsien 1990, Sola et al. 2004) of this quantal analysis (Figure 5K) is consistent with the interpretation that consecutive activation of synaptic connections between L5 PNs in the PPC leads to a reduction in active release sites (*n*) specifically in aRMS exposed mice.

To test whether the postsynaptic composition of PN-to-PN synapses was impacted by aRMS, we employed peak-scaled nonstationary fluctuation analysis (Sigworth 1980, Yu et al. 2016, Traynelis, Silver, and Cull-Candy 1993). To estimate single-channel current and the number of channels per synapse, we fitted the mean-variance plot using a least-squares algorithm (Figure 5L, see Methods). We found that neither the mean single-channel current (control: 1.21 ± 0.43 pA, n = 15, aRMS: 1.84 ± 0.65, n = 12, p = 0.1, Welch’s t-test, Figure 5M) nor the estimated number of channels per synapse (control: 10.74 ± 4.5, n = 15, aRMS: 6.71 ± 1.8, n = 12, p = 0.09, Welch’s t-test, Figure 5N) was different between control and aRMS animals.

Overall, our paired patch clamp recordings provided further evidence that the connection probability between pyramidal neurons in the local PPC network remains unaffected by aRMS. However, we observed a stress-induced reduction in the decay phase of unitary response currents and a reduction in the number of active release sites with consecutive activation between connected pyramidal neurons.

### aRMS increases GABAergic tone in the PPC

Cortical activity is sculpted by the action of inhibitory interneurons (Garcia Del Molino et al. 2017, Compte et al. 2000, Najafi et al. 2020, Papoutsi, Sidiropoulou, and Poirazi 2014, Sadeh and Clopath 2021). Indeed, our computational model predicted that inhibitory synaptic conductances mediated by GABAa-, and GABAb-receptors play a substantial role in persistent firing. To test the effect of aRMS on inhibition in the PPC, we measured inhibitory responses in L5 PNs evoked by electrical stimulation. First, we utilized a paired-pulse stimulus protocol and pharmacologically isolated inhibitory postsynaptic potentials (IPSPs) mediated by GABAa-receptors (CGP55845, 1μM, Figure 6A). To determine the effect of aRMS on response magnitude, we gradually increased stimulus intensity until the IPSP amplitudes saturated. While larger stimuli evoked larger responses (fixed effect of stimulus intensity: F_1.968, 64.30_ = 8.859, p = 0.0004, two-way mixed effects model applied to the first peak, n_ctr_ = 21, n_aRMS_ = 16, Figure 6B), we detected no effect of aRMS on the magnitude of the first response peak (fixed effect of stress: F_1, 35_ = 0.6742, p = 0.42). However, we found that aRMS markedly impacted the short-term dynamics of GABAa-mediated responses. Responses to the second stimulus were smaller than to the first one in controls (fixed effect of first vs second peak: F_1, 30_ = 5.879, p = 0.0216). In contrast, we did not see such short-term depression in aRMS mice (F_1, 40_ = 0.05497, p = 0.8158). Indeed, a three-way mixed-effects model showed significant interaction between first and second peak amplitudes and the stress condition (F_1, 35_ = 14.91, p = 0.0005, Figure 6B). Reinforcing this finding was the marked stress effect on the paired pulse ratio of GABAa responses (F_1, 35_ = 25.43, p < 0.001, two-way mixed effects model, Figure 6C).

**Figure 6:**
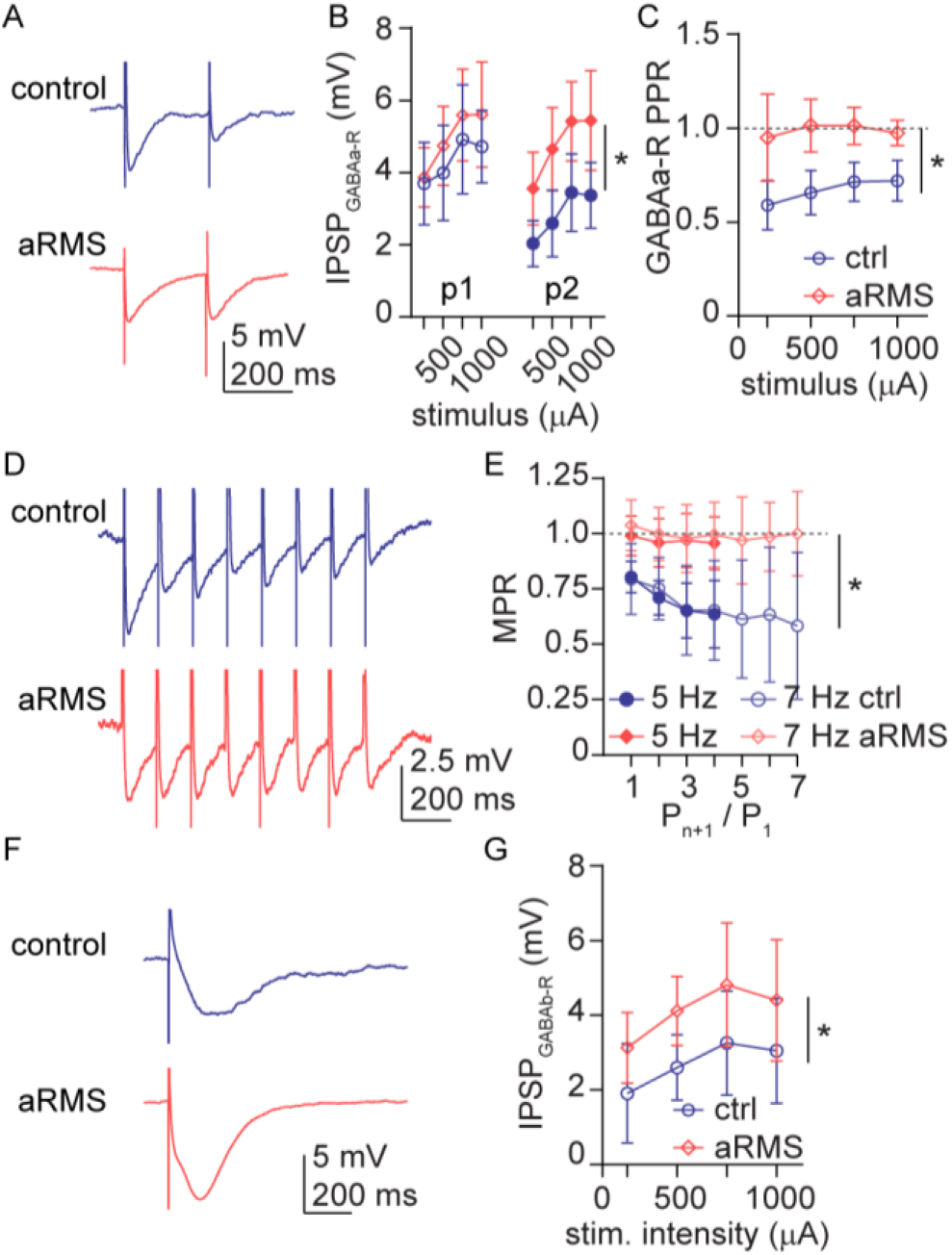
aRMS increases GABAergic tone in the PPC. (**A**) Representative traces illustrating evoked GABAa receptor mediated currents in control (blue) and aRMS exposed (salmon) mice. (**B**) First (p1) and second (p2) GABAa mediated IPSP peak amplitudes in response to increasing stimulus magnitudes. (**C**) Paired pulse ratio (p2 / p1) of GABAa receptor currents at different stimulus strengths. (**D**) Example traces showing GABAa mediated responses to a 7 Hz stimulus train in control (blue) and aRMS (salmon) mice. (**E**) Multi-pulse ratio in response to 5 Hz (solid markers) or 7 Hz (open markers) stimulus trains in control (blue) and aRMS (salmon) mice. (**F**) Example traces of GABAb receptor mediated IPSPs in control (blue) and aRMS (salmon) animals. (**G**) GABAb-mediated IPSP peak amplitudes in response to increasing stimulus strength in control (blue) and aRMS (salmon) mice. Markers indicate mean, error bars show ± 95 CI. *: p < 0.05.

The typical frequency of persistent firing in L5 PNs was between 5 and 7 Hz. To test whether aRMS could impact short-term GABAa dynamics at these frequencies, we collected IPSP data at either 5 or 7 Hz stimulus rates (Figure 6D) and displayed the relationship of the first peak to subsequent responses as a multi-pulse ratio. As with paired pulse stimuli above, we found that inhibitory responses were depressing in controls but remained consistent in magnitude in aRMS mice (main effect of stress at 5Hz: F_1, 17_ = 14.56, p = 0.0014, n_ctr_ = 12, n_aRMS_ = 7; main effect of stress at 7Hz: F_1, 12_ = 9.969, p = 0.0083, n_ctr_ = 8, n_aRMS_ = 6, two-way ANOVA, Figure 6E).

Next, we pharmacologically isolated inhibitory responses mediated by GABAb-receptors (PTX, 50μM Figure 6F). To determine the effect of aRMS on the magnitude of GABAb dependent inhibition, we recorded responses at increasing stimulus strength. As expected, larger stimuli evoked larger responses in both populations (fixed effect of stimulus intensity: F_2.111, 23.92_ = 6.828, p = 0.004011, n_ctr_ = 8, n_aRMS_ = 6, two-way mixed effects model). In addition, we found a consistent increase in response magnitude in aRMS mice (fixed effect of stress: F_1, 12_ = 6.080, p = 0.03, Figure 6G).

These data suggest that aRMS exposure enhances GABAergic inhibition in the PPC both by reducing short-term depression of GABAa responses and increasing the magnitude of GABAb mediated inhibition.

### A combination of factors is necessary to explain the aRMS-induced reduction in persistent firing in the PPC

Our data suggests that aRMS altered a variety of physiological attributes of the recurrent circuit in PPC layer 5. These include the cell intrinsic input resistance and properties of the excitatory and inhibitory connectivity within the local network. To determine which of these factors may be responsible for reducing sustained action potential firing in aRMS exposed mice, we turned back to our computational framework. In the same network model we used to make predictions about potential mechanisms, we set external stimulus duration to 2 seconds and the properties of the external drive to produce persistent firing with a 100% probability (NMDA weight set to 0.43, see Methods). To remain faithful to our physiological observations, we ran a “control” model where the decay time of NMDA-receptors in the external synaptic drive was set to 90 ms, and an “aRMS” model where NMDA decay was set to 150ms. Keeping all other parameters fixed, we systematically adjusted the physiological attributes we knew to be impacted by aRMS to values similar to what we found in our experimental data. Independently changing input resistance, the short-term dynamics of GABAa responses, the magnitude of GABAb responses, the synaptic decay, or short-term depression of recurrent connections, we found that each parameter adjusted alone had no effect on persistent firing probability in the stressed network (Figure 7A). Depending on the specific combination, the effect of altering two parameters could be negated by the increased NMDA-current decay in the stressed network, or it could have a small (10 to 30%) effect on persistent firing. Simultaneous inclusion of three parameters enhanced the effect to 10 to 60% reduction while including four or all five parameters resulted in a substantial 50 to 90% reduction of persistent firing. This effect was commensurable with the approximately 66% reduction of persistent firing in our *ex vivo* experiments. Any combination of parameters had a consistently larger impact on the control network due to the longer NMDA-receptor decay in aRMS mice – a likely compensatory mechanism following stress exposure (Figure 7A).

**Figure 7:**
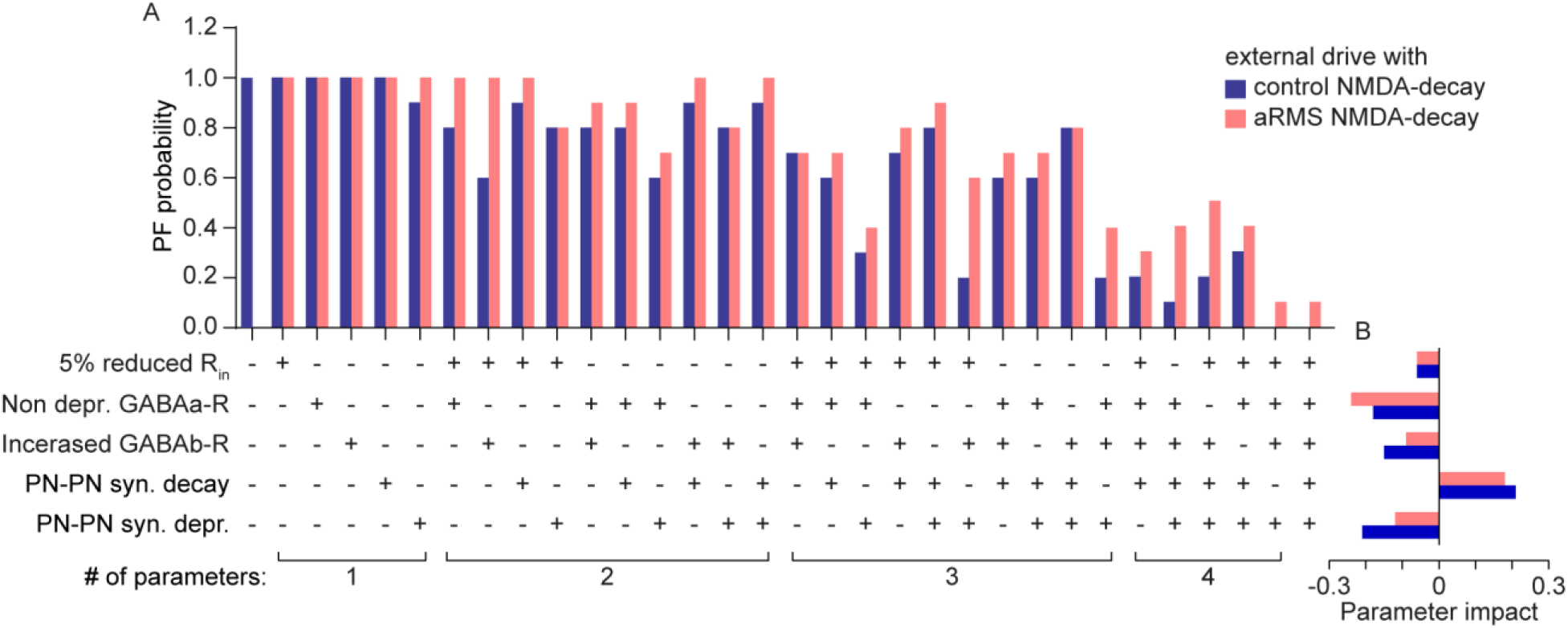
A combination of aRMS-induced changes is necessary to reduce persistent firing. (**A**) Bars indicate persistent firing probability in a computational model simulating the presence (+) or absence (−) of an aRMS-induced change in key physiological parameters. The external synaptic drive to the network was implemented as observed in recordings from control (blue) or in aRMS exposed (salmon) animals. The number of parameters altered simultaneously is indicated at the bottom. (**B**) The histogram indicates the estimated impact of each parameter on persistent firing probability.

To estimate the impact of each aRMS-induced physiological change in our model, we calculated the difference between the persistent firing probability with the parameter set to the control value from the probability with the parameter set to aRMS value. A more negative value indicates a stronger reduction in persistent firing when the parameter is set to aRMS. We found that the short-term dynamics of GABAa responses had the largest impact on persistent firing, followed by the short-term dynamics of recurrent synapses and the magnitude of GABAb responses (Figure 7B).

## Discussion

In this study, we examined how repeated exposure to multiple concurrent stresses in adolescence (aRMS) affected persistent firing in layer 5 pyramidal neurons (L5 PNs) of the posterior parietal cortex (PPC). We found that aRMS significantly decreased both the duration and frequency of persistent firing. Using a combination of computational modeling, viral tracing, and patch-clamp electrophysiology, we systematically analyzed intrinsic properties, local recurrent connectivity, as well as excitatory and inhibitory synaptic dynamics to identify mechanisms underpinning this deficit. While intrinsic excitability was mostly preserved, aRMS decreased the input resistance of L5 PNs. Although local recurrent connectivity remained unchanged, we found a notable reduction in the decay time constant and an increase in short-term depression at PN-PN excitatory synapses. In addition, we observed heightened inhibitory tone via actions through both fast GABAa- and slower GABAb-receptors following aRMS. Incorporating these changes into our network model allowed us to test the impact of aRMS-induced physiological changes. We found that changing at least two factors was necessary to impact persistent firing, the effect gradually increasing with more parameters altered. Specifically, the reorganization of inhibitory dynamics and the short-term depression of recurrent excitatory synapses had the largest impact on persistent firing. These findings suggest that aRMS impedes persistent firing through a combination of orthogonal mechanisms rather than a single dominant causal factor.

### Minimal effect of aRMS on intrinsic properties of PNs in the PPC

Chronic stress has been shown to have diverse effects on the excitability and intrinsic properties of cortical pyramidal neurons (Urban and Valentino 2017, Song et al. 2019, Anderson et al. 2019). Our whole-cell patch-clamp recordings revealed no changes in action potential threshold, sag amplitude, or the overall input-output relationship of L5 PNs, while input resistance was slightly decreased in PNs from aRMS mice. A decrease in input resistance can reduce the depolarizing impact of excitatory postsynaptic currents, thereby diminishing the ability of recurrent inputs to maintain persistent firing, an effect captured in our predictive model. Similarly, stress-induced reduction in input resistance has been previously observed by us and others across various stress models (Nawreen, Baccei, and Herman 2021, Kole, Czeh, and Fuchs 2004, Libovner et al. 2020). Backtesting our physiology results in the network model *in silico* indicated that while input resistance alone produced a small reduction in persistent firing probability, it was likely insufficient to account for the diminished persistent firing we observed, suggesting that it acted synergistically with other, synaptic changes.

### aRMS only impacts the NMDA component of overall synaptic drive in the PPC

Chronic exposure to stress is well-known to alter excitatory synapses and glutamatergic transmission in the neocortex (Bueno-Fernandez et al. 2021, Musazzi, Treccani, and Popoli 2015, Popoli et al. 2011, Rodrigues et al. 2024, McKlveen et al. 2016). Here, we examined the composition of the excitatory synaptic drive in the PPC. We found that NMDA receptor-mediated currents in aRMS mice exhibited slower decay, whereas AMPA receptor-mediated responses and the ratio of the two receptor types remained unaffected. Numerous previous studies found increased NMDA receptor expression or function following repeated stress exposure, although the effects are often brain region specific (Zhou et al. 2018, Sun et al. 2020, Lei and Tejani-Butt 2010). Prolonged NMDA decay could facilitate temporal integration and enhance network stability (Compte et al. 2000, Wang 1999), a result reflected in our predictive model as well. Thus, increased NMDA receptor decay contrasts with our observation of stress-induced reduction in persistent firing. An important caveat in these experiments comes from the nature of the stimulus: bipolar electrodes placed in the cortex are non-specific, activating both local synapses as well as long-range inputs. As our later results demonstrate, there is a disconnect between the effect of aRMS on overall excitatory drive and PN-to-PN recurrent synapses. We speculate that the increased NMDA receptor time constant is a compensatory mechanism that is ultimately outweighed by other stress-induced synaptic changes in the local network. To demonstrate this, we included the longer NMDA receptor decay in our *in silico* backtesting, creating separate “control” and “stressed” model conditions. Our results show that the effects of stress-altered physiological parameters are reduced in the “stressed” model but are not fully compensated.

### The number of recurrent connections are preserved in aRMS mice, but synaptic properties are altered

Robust evidence suggests that repeated stress exposure eliminates excitatory synapses from the neocortex (Woo et al. 2021, Csabai, Wiborg, and Czeh 2018, Radley et al. 2006). Based on this, a reasonable hypothesis was that aRMS dampens persistent activity by reducing the excitatory connectivity within the recurrent cortical network (Wang 1999, Hass et al. 2022, Barak and Tsodyks 2007, Papoutsi, Sidiropoulou, and Poirazi 2014). In agreement, our predictive model indicates that either reducing the number of PN-to-PN connections or the synaptic strength between the nodes of the network will impede persistent firing. However, rabies virus-based trans-synaptic tracing and paired whole cell recordings showed no change in the number of recurrent connections between L5 PNs in the PPC. These data suggest that the aRMS-induced reduction in persistent firing is not due to loss of excitatory connections within the recurrent circuit. Conversely, our paired recordings revealed functional changes in recurrent synapses. The faster EPSP decay kinetics we measured after aRMS would reduce the integration window and act to decrease persistent firing (Lei and Tejani-Butt 2010, Sidiropoulou and Poirazi 2012, Sun et al. 2020, Zhou et al. 2018). This stress-induced change in decay kinetics could reflect an increase in the proportion of the NR2A receptor subunits in recurrent excitatory synapses (Lindlbauer et al. 1998, Santucci and Raghavachari 2008). In addition, our paired pulse ratio measurements showed that following repeated activation, recurrent excitatory synapses in aRMS mice became less efficient. Quantal analysis indicated that the difference was likely due to the reduction of active release sites in stressed animals (Kloc and Maffei 2014, Sola et al. 2004, Wang, Fontanini, and Maffei 2012, Wernig 1975). In contrast, we did not find stress effects on the postsynaptic side. These changes would reduce the reliability of the excitatory drive in the recurrent network. Indeed, when we backtested our findings *in silico*, we found that faster synaptic decay and increased short-term depression can alter the ability of the recurrent network to produce persistent firing. Notably, the observed stress-effects were specific to PN-to-PN synapses captured via paired whole cell recordings and were not obvious from non-specific electrical stimulation.

### Increased GABAergic inhibition following aRMS

The effect of chronic stress on cortical inhibition is somewhat controversial: some authors found enhanced inhibition in the prefrontal cortex (McKlveen et al. 2016), while others observed the opposite effect (Czeh et al. 2018). Notably, the above studies used similar aged (adolescent to young adult) rats and the same chronic variable stress paradigm, with the main difference being exposure time. Results from adolescent mice showed increased inhibition in the prefrontal cortex somewhat more consistently (Anderson et al. 2019, Rodrigues et al. 2024). We observed two avenues of enhanced GABAergic tone following aRMS. First, GABAa receptor-mediated responses lost their typical short-term depression. This change in short-term kinetics may be due to altered presynaptic GABA release (Regehr 2012) or reuptake mechanisms (Ma et al. 2016, Overstreet and Westbrook 2003). In addition, GABAb receptor-mediated responses showed increased magnitude. Both effects could suppress the sustained firing ability of L5 PNs. Non-depressing GABAa responses could prevent excitatory build-up during repetitive activity, while stronger GABAb-mediated inhibition would provide slow hyperpolarization, limiting the maintenance of persistent activity. These findings align with our own computational predictions and prior reports showing that stress can enhance inhibitory reliability in cortical circuits (Compte et al. 2000, McKlveen et al. 2016). Our *in silico* backtesting demonstrated that these inhibitory changes had some of the largest impact on persistent firing probability, indicating a central role in stress-induced network disruption.

### Multiplexing aRMS-induced changes is necessary to reduce persistent firing

We integrated the above physiological parameters into a computational model to determine which factors may be necessary and sufficient to reduce persistent firing, as we observed in stressed mice. Our model suggests that no single parameter could account for the observed reduction in persistent firing on its own. Instead, a combination of two or more aRMS-induced changes, particularly increased inhibitory tone, weakened glutamate release over repetitive stimulus, and reduced input resistance, acted synergistically to suppress the probability of persistent firing in the model. This may not be too surprising given the large number of factors that can influence persistent firing both at the cellular and network levels (Murray, Jaramillo, and Wang 2017, Masse et al. 2019, Zylberberg and Strowbridge 2017, Lin et al. 2020, Barak and Tsodyks 2007, Sidiropoulou and Poirazi 2012). While the slower NMDA decay of overall excitatory drive observed in aRMS mice moved the model towards more persistent firing (Compte et al. 2000, Wang 1999), it could not fully compensate for stress-induced changes in most cases. Our results indicate that repeated stress exposure produces a multitude of converging physiological changes. This distributed mechanism may explain why stress-induced cognitive deficits present such a challenging therapeutic.

### Therapeutic Implications

Chronic stress exposure has a devastating effect on cognitive function and is considered one of the key risk factors for many mood and anxiety disorders (Girotti, Bulin, and Carreno 2024, McEwen 2017, McEwen, Nasca, and Gray 2016, Miguel-Hidalgo 2013). Despite its prevalence, effective pharmacological or non-pharmacological treatment options for stress-induced cognitive impairment are lacking (Colwell et al. 2022, Girotti, Bulin, and Carreno 2024). A multitude of the impacted cognitive processes rely on the ability of the cortex to produce sustained activity patterns – a function we have shown to be disrupted after stress exposure. Our results indicate that GABAergic signaling may be a potent therapeutic target for restoring persistent firing and thus improving cognitive outcomes following stress exposure. There are a few studies supporting this notion. In chronically stressed rats, the GABAa receptor antagonist bicuculline improved hippocampus-dependent spatial memory, though stressed animals required higher doses than controls (Nishimura, Ortiz, and Conrad 2017). In human subjects, the benzodiazepine-site antagonist flumazenil enhanced working memory in both schizophrenia patients and healthy controls, while the GABAa receptor agonist lorazepam impaired working memory (Menzies et al. 2007). More selective compounds, such as α5-subunit GABAa receptor negative allosteric modulators and GABAb receptor antagonists, have been shown in stress and depression models to reduce inhibitory tone, increase glutamatergic transmission, and improve prefrontal cortex-dependent cognitive performance (Maguire 2014, Walton et al. 2023, Hipp et al. 2021). Ketamine, although primarily an NMDA receptor antagonist, can also act through disinhibition by blocking NMDA receptors on inhibitory interneurons, specifically in the parvalbumin subtype, which transiently reduces GABAergic drive, enhances pyramidal cell activity, and promotes synaptic plasticity (Widman and McMahon 2018). These findings, together with our data, suggest that pharmacological strategies that curb stress-enhanced inhibition either directly through GABA receptor modulation or indirectly through disinhibitory mechanisms may help restore persistent firing and associated cognitive functions.

## Acknowledgements

We thank Dr. Lin C. Cheng for his assistance with data visualization and Mikko Oijala for his contribution to our stress setup and other engineering efforts.

## Methods

### Animals

All experiments were performed in accordance with the NIH guidelines on laboratory animal welfare and were approved by the University of California, Irvine Institutional Animal Care and Use Committee. Male mice were group housed in a quiet, uncrowded facility on a 12 h light / dark cycle, with *ad libitum* access to food and water. For all procedures, male C57BL/6 mice were either purchased from Charles River or bred in-house. Stress and control cohorts were counterbalanced within litters and shipped batches.

### Stress Paradigm

Male mice were randomly assigned to either control or stress groups. Control mice remained in their home cages. Mice in the stress group were subjected to the previously described repeated exposure to multiple concurrent stressors paradigm during early-to mid-adolescence (postnatal day (P)30-40, termed aRMS for short) (Libovner et al. 2020, Fariborzi et al. 2021, Hokenson et al. 2020). Briefly, each mouse was restrained in a well vented 50ml conical tube, and 6-8 mice were placed on a laboratory rocker. Here, animals were simultaneously subjected to bright light, jostling, and loud noises (15-30 kHz at 0.5-3 s random intervals) for 1 hour per day for 10 consecutive days. At P40 (approximately 1 day after the end of the stress period), age-matched controls and aRMS mice were sacrificed for acute slice electrophysiology.

### Slice electrophysiology

For electrophysiology recordings, 400μm thick coronal brain slices containing the PPC were prepared on a vibrating microtome (smz7000-2, Campden Instruments). Immediately after sectioning, slices were maintained at 32°C for 15 minutes in solution comprised of (in mM): 110 choline, 25 NaHCO_3_, 1.25 NaH_2_PO_4_, 3 KCl, 7 MgCl_2_, 0.5 CaCl_2_, 10 glucose, 11.6 sodium ascorbate, and 3.1 sodium pyruvate, and bubbled with 95% O_2_ and 5% CO_2_. Slices were then allowed to recover for at least 20 minutes in oxygenated room-temperature aCSF containing (in mM): 126 NaCl, 25 NaHCO_3_, 10 Glucose, 3 KCl, 2 CaCl_2_, 1.25 NaH_2_PO_4_, and 1 MgSO_4_ prior to recording (Libovner et al. 2020, Rindner, Proddutur, and Lur 2022). Recording aCSF for persistent firing experiments was modified to contain 3.5mM KCl, 1.2mM CaCl2, and 10µM carbochol, a cholinergic (both muscarinic and nicotinic receptors) agonist to mimic elevated acetylcholine tone during awake conditions and improve the probability of evoking persistent activity (Major, Vijayraghavan, and Everling 2018, McCormick and Contreras 2001, Vijayraghavan, Major, and Everling 2018, Zhou et al. 2011).

All experiments were conducted close to physiological temperature (32–34 °C) in a submersion type recording chamber mounted on an Olympus BX51-WI microscope. The hippocampus was used as a histological landmark to identify PPC in coronal sections. Whole-cell current and voltage clamp recordings were performed in layer 5 pyramidal neurons in the mediolateral section of PPC between 1600–1800 μm from the midline and 400-600 µm from the pia surface to minimize the influence of potential subregion-specific differences. Cells were identified with video-infrared / differential interference contrast microscopy using a 60X water-immersion objective.

For all current clamp recordings, glass electrodes (2–4 MΩ) were filled with internal solution containing the following (in mM): 135 KMeSO3, 10 HEPES, 4 MgCl2, 4 Na2ATP, 0.4 NaGTP, and 10 sodium creatine phosphate, and adjusted to pH 7.3 with KOH. For voltage clamp recordings of evoked AMPA and NMDA currents, internal solution containing cesium was used for better space clamp (in mM): 135 CsMeSO3, 10 HEPES, 4 MgCl2, 4 Na2ATP, 0.4 NaGTP, and 10 sodium creatine phosphate, and adjusted to pH 7.3 with CsOH. Series resistance was 10– 25 MΩ and bridge compensated in current clamp. Electrophysiological recordings were made using a MultiClamp 700B Amplifier (Molecular Devices), filtered at 4 kHz, and digitized at 10 kHz. Data was acquired using National Instruments DAQ boards and Wavesurfer software written in MATLAB (Howard Hughes Medical Institute Janelia Research Campus). Offline analysis was performed using Clampfit 10.7 and custom routines written in MATLAB or Python 3.7 (Anaconda distribution).

To examine the active and passive intrinsic properties of L5 PNs, we recorded membrane potential responses to a series of 0.5 second long square pulse current injections, ranging from - 350 to 400 pA with a step size of 50 pA.

### Electrical stimulation

Postsynaptic AMPA- and NMDA receptor-mediated excitatory postsynaptic currents (EPSCs) were recorded with membrane voltage clamped at −70 mV and +40 mV, respectively. A bipolar stimulating electrode was placed approximately 100–150 μm from the recorded L5 PN and used to evoke postsynaptic responses with a stimulus current of 50–250 μA for 500 μs. Inhibitory currents through GABAa and GABAb receptors were blocked (PTX, 50 μM, and CGP55845, 1 μM) to pharmacologically isolate excitatory responses. In a subset of recordings we confirmed AMPA or NMDA responses via bath application of NBQX (10 μM) or AP5 (50 μM).

Postsynaptic GABAa- and GABAb receptor-mediated inhibitory postsynaptic potentials (IPSPs) were recorded in current clamp, with the membrane potential held at −70mV in the presence of glutamate receptor blockers NBQX (10 μM) and AP5 (50 μM). Responses mediated by either GABAa or GABAb receptors were pharmacologically isolated using CGP55845 (1 μM), or picrotoxin (100 μM), respectively. The bipolar stimulation electrode was placed approximately 100–150 μm from the recorded L5 PN to evoke inhibitory responses with increasing stimulus currents ranging from 250 μA to 1 mA for 500 μs. When recording GABAa responses we used a paired pulse stimulation protocol with an interstimulus interval of 250 ms or a pulse-train of five stimuli at 5 or 7 Hz frequency.

For all stimulus maps we collected and averaged 10 repeats with an interval of 10 seconds.

### Paired recordings

Dual whole cell recordings were conducted by patching two nearby (within 150 µm) layer 5 PNs simultaneously. Connectivity between cells was examined by holding one (presynaptic) PN in current clamp and the other (postsynaptic) cell in voltage clamp at −70 mV. We elicited 10 action potentials in the presynaptic PN via a square pulse train of current injections (0.5s total duration, 20Hz frequency, 2ms pulses at an amplitude of 250 pA) and recorded excitatory postsynaptic currents (EPSCs) in the postsynaptic PN. We collected 5-10 repeats with an interval of 10 seconds. Each pair of cells were tested for monosynaptic or reciprocal connections by switching between the pre-, and postsynaptic roles.

### Quantal analysis

We examined the properties of excitatory transmission in recurrent synapses using quantal analysis. To estimate quantal properties, we took advantage of synaptically coupled paired recordings (Sola et al. 2004, van Huijstee and Kessels 2020). We measured the peak amplitude of the first and the tenth EPSCs across a minimum of 5 repeats and calculated the mean amplitude (M) and standard deviation (`σ`) of the responses. We determined the variance-to-mean ratio (VMR) as

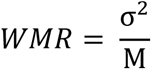

WMR is thought to be dependent on presynaptic release probability (P) and postsynaptic response to the release of a single vesicle or quantal size (Q), but not on the number of active release sites (N) (Lupica, Proctor, and Dunwiddie 1992, van Huijstee and Kessels 2020). We also calculated the inverse square of the coefficient of variation (1/CV^2^) as

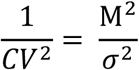

This metric depends on N and P, but is independent of Q (Malinow and Tsien 1990, van Huijstee and Kessels 2020). The above measures are based on a binomial model of synaptic transmission first applied to the neuromuscular junction (McLachlan 1978, Wernig 1975). This model allows us to interpret the inequality:

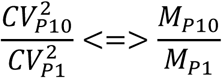

where P1 refers to the first, and P10 to the tenth response (Kloc and Maffei 2014, Sola et al. 2004). This inequality can be visually represented in a topographical plot that provides insights into changes in neurotransmission (Bekkers and Stevens 1990, Malinow and Tsien 1990). The model is frequently interpreted as follows: CV_P10_^2^/ CV_P1_ ^2^ > M_P10_/M_P1_, indicates that both N and P changed; CV_P10_^2^/ CV_P1_ ^2^ = M_P10_/M_P1_, suggests that only N changed; CV_P10_^2^/ CV_P1_ ^2^ < M_P10_/M_P1_, indicates that neither N nor P changed, suggesting an exclusive effect on Q.

### Nonstationary fluctuation analysis

Nonstationary variance analysis (Proddutur et al. 2023, Sigworth 1980, Yu et al. 2016) was performed on postsynaptic current responses evoked by presynaptic action potentials in synaptically coupled L5 PNs. Stable recordings were pooled from individual groups to isolate fluctuations in the current decay attributable to stochastic channel gating, the mean waveform was scaled to the peak of individual EPSCs (Traynelis, Silver, and Cull-Candy 1993, De Koninck and Mody 1994). Successful responses were used to calculate the ensemble mean current (I) and peak-scaled variance (σPS2) for each data point. Plots of variance versus current were fit with the equation:

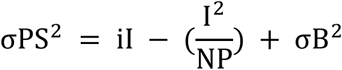

where i is the weighted-mean single-channel current, NP is the number of channels open at peak synaptic current, and σB^2^ is the background variance (Brickley, Cull-Candy, and Farrant 1999). Using the average of the first evoked response, we calculated the peak amplitude as the maximum dip from the baseline. Rise time was measured as the time from 10% to 90% of the peak during the rising phase, and decay time as the time from 90% to 10% of the peak during the decay phase.

### Rabies based trans-synaptic tracing

To access the local network of layer 5 pyramidal neurons in the PPC, eight days before aRMS (at p22) we expressed Cre-recombinase in L5 PNs by injecting 300 nl retrograde adeno-associated virus (rgAAV.hSyn.Cre) to the dorsal striatum ipsilateral to our target hemisphere (coordinates in mm from the Bregma: AP:-0.7, ML:1.5, DV:2.4). In addition, we injected 400 nl of Cre-dependent rabies helper constructs (a 1:1 mix of AAV.Flex^FRT^.TVA66T.mCherry and AAV.Flex^FRT^.RABV-G) in the PPC (coordinates in mm from the Bregma: AP −2.0, ML:1.6, DV: 0.5). The day after the final aRMS session (p41), we transduced the PPC with EnvA-pseudotyped rabies virus (RABV). In a subset of mice, we omitted hSyn.Cre retrograde AAV, but injected both helper constructs and EnvA-pseudotyped rabies virus (RABV) as described above. In the absence of Cre expression, we did not see any non-specific helper virus or RABV expression in the slices (data not shown). Animals were sacrificed after 7 days of RABV expression via transcardial perfusion of phosphate buffered saline (PBS) followed by 4% paraformaldehyde (PFA) dissolved in PBS. Brains were post-fixed in 4% PFA overnight at 4°C prior to cutting 50 µm thick sections on a vibrating blade microtome (Compresstome). PPC containing sections were mounted on microscope slides and sealed under a cover slip with Prolong Diamond antifade reagent.

### Imaging and cell counting

Histological sections from the rabies virus-based trans-synaptic tracing experiments were visualized through a 10x (0.3 NA) objective on a Zeiss Axio Observer Z1 inverted fluorescence microscope connected to a Hammamatsu ORCA-ER monochrome CCD camera. Fluorescence excitation light was provided by an LED light source, and emissions were collected through appropriate FITC and Texas Red filter sets. To avoid bias, all histological sections were imaged at the same exposure settings with the experimenter blinded to the group condition. We used three representative PPC slices per mouse in each group.

To quantify starter cells and presynaptic neurons, we counted fluorescent cells using automated batch processing in CellProfiler (Lamprecht, Sabatini, and Carpenter 2007, McQuin et al. 2018, Libovner et al. 2020). Our pipeline included an object identification algorithm with object size ranging from 5 to 50 pixels in the mCherry (Cre and Helper AAV positive cells) and GFP (Presynaptic neurons) channels. To identify cells with colocalization, we used rank-weighted colocalization and overlap coefficients metrics to identify starter neurons (postsynaptic cells co-expressing mCherry and GFP).

### Computational modeling

We adapted a previously published persistent firing network model (Papoutsi, Sidiropoulou, and Poirazi 2014) implemented in the NEURON simulation environment (Hines and Carnevale 1997). The network includes 7 pyramidal neurons and 2 interneurons. The morphology of pyramidal cells included a soma, axon, basal dendrite, proximal apical dendrite, and distal apical dendrite, while interneurons had a simplified morphology that only included somatic and axonal compartments. Active and passive channel parameters for both cell types were set to the original published model. Similarly, synaptic kinetics were adapted from the original model, where NMDA : AMPA ratios were adjusted by increasing NMDA conductance while keeping AMPA constant. To keep terminology consistent with the original model, an NMDA weight of 0.35, or 0.43 produced an iNMDA : iAMPA ratio of 1.5, or 1.8, respectively. Persistent firing in the network model was generated by an external synaptic input using 10 to 40 pulses at 20Hz applied to 90 excitatory synapses on proximal dendrites of all PNs. Additionally, to mimic *in vivo*-like activity, all neurons in the microcircuit received Poisson-distributed current injection and background input via 120 glutamatergic synapses on PNs and 40 synapses on INs, which were randomly distributed across dendritic locations and activated by a 6 Hz Poisson train with −250 to +50 ms jitter, producing correlated subthreshold fluctuations and spontaneous firing rates comparable to *in vivo* quiet wakefulness.

We specifically modified and implemented the GABAa receptor-mediated synaptic mechanism at IN to PN synapses to include short-term depression with a fast decay component set to 0.5 and a recovery time constant of 380 ms (Varela et al. 1997). Similarly, the short-term depression of PN-PN synapses were implemented with a depression factor set to 0.5 and a recovery time constant of 250 ms. This effectively scaled the synaptic strength on each presynaptic spike by a multiplicative factor of 0.5. We implemented a reduction in PN-PN synaptic decay by increasing the AMPA receptor unbinding rate from the original value of ‘0.15/ ms’ to a faster closing rate of ‘0.35 / ms’, which accelerated the closing kinetics of the synapse and results in a shorter EPSC duration. To implement a reduction in the input resistance of PN, we modified the passive conductance ‘g_pas’ mechanism in all the compartments of PN model neurons.

### Data analysis and statistics

All analyses were performed by an experimenter blinded to treatment groups, followed by unblinding to categorize treatment groups for statistical analysis. Electrophysiology recordings were discontinued if series resistance increased by more than 20% or if the resting membrane potential was less than −54mV. Cells were also excluded from analysis if they were lost before or after drug flow in, or if fewer than 3 sweeps were recorded. To measure the intrinsic properties of L5 PNs, action potential peaks were automatically identified using a peak finder algorithm, that computed the number, height, timing, and width of action potentials. Recorded neurons were initially held at −70 mV, and responses to 0.5 second positive and negative current injections were examined to determine active and passive properties. Input resistance was measured by observing the membrane response to a +50 pA current injection, and SAG was measured at a −250 pA current step as described in (Rindner, Proddutur, and Lur 2022). The threshold of the action potential was determined by calculating the first derivative (dV / dt) of the voltage trace during the first action potential and setting 10 mV / ms as the threshold level for action potential initiation (Gupta et al. 2012).

We calculated persistent firing probability in our computational model by averaging the probability recorded from the seven pyramidal cells across 10 randomized simulations. For each cell in the model, after the network stimulus drive has ended, we determined the persistent firing probability as a success (1) if the duration lasted 10 seconds or more, or as a failure (0) if it lasted less than 10 seconds.

Statistical analyses were conducted in Prism (GraphPad). Single variables were compared using t-tests when populations had similar standard deviation. When the standard deviations were substantially different, we used t-tests with Welch’s correction, we refer to this as Welch’s t-test in the body of the text. To compare multiple variables, we used the appropriate two-, or three-way ANOVA models where possible, and substituted with mixed effects models where necessary due to missing values. All data in the text and figures are displayed as mean ± 95% confidence intervals (CI) with individual data points showcased when appropriate.

