## Supplementary Figures for "Multiplexed changes in synaptic transmission underlie stress-induced reduction of persistent firing in the parietal cortex"

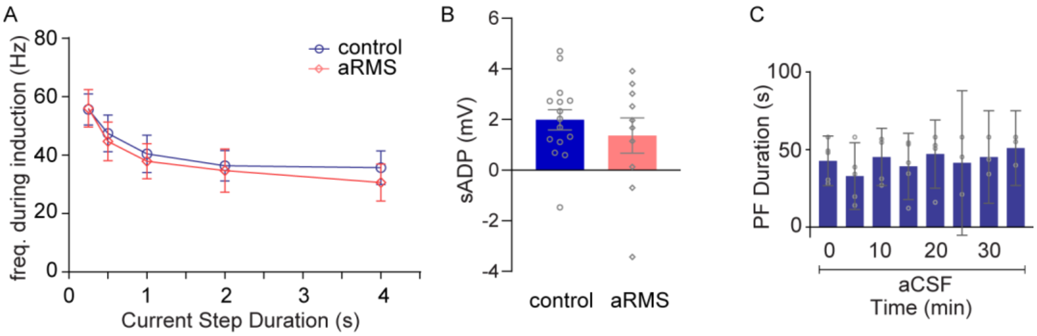


**Supplementary Figure 1:** (**A**) Frequency of action potential firing in control (blue) vs aRMS (salmon) mice during the various duration current injections (250pA) used for inducing persistent firing. (**B**) Amplitude of the slow after depolarization (sADP) following the induction protocol applied to control (blue) or aRMS (salmon) cells held at -75mV in current clamp. (**C**) Persistent firing duration remains stable over time and through repeated inductions in aCSF. Bars represent population mean ± 95% CI.

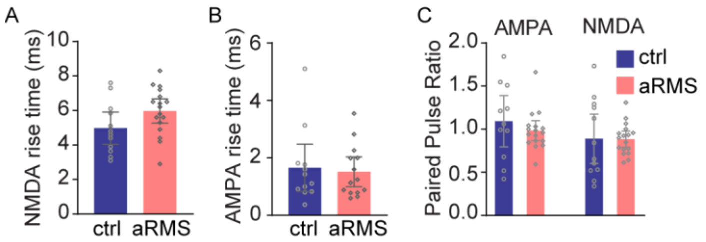


**Supplementary Figure 2:** (**A**) Rise time of electrical stimulation evoked NMDA receptor responses in control (blue) and aRMS (salmon) mice. (**B**) Rise time of electrical stimulation evoked AMPA receptor mediated responses. (**C**) Pair pulse ratio of electrical stimulation evoked AMPA-, and NMDA receptor mediated responses in control (blue) and aRMS (salmon) mice. Bars represent population mean ± 95 CI.

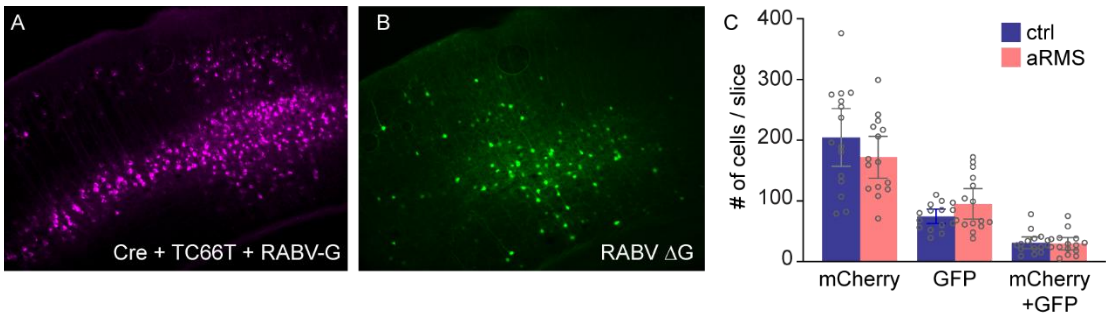


**Supplementary Figure 3:** (**A**) Representative micrograph showing helper virus (mCherry fluorescent tag) expression in the PPC (mCherry pseudocolored as magenta). (**B**) Representative image showing RABVΔG expression (GFP fluorescence) in presynaptic neurons in the PPC (green). (**C**) The number of mCherry positive (helper construct expressing), GFP positive (presynaptic cells), and mCherry + GFP positive neurons (starter, postsynaptic neurons) in control (blue) and aRMS (salmon) mice. Bars represent population mean ± 95 CI.
